# Heteroplasmy variability in individuals with biparentally inherited mitochondrial DNA

**DOI:** 10.1101/2020.02.26.939405

**Authors:** Jesse Slone, Weiwei Zou, Shiyu Luo, Eric S Schmitt, Stella Maris Chen, Xinjian Wang, Jenice Brown, Meghan Bromwell, Yin-Hsiu Chien, Wuh-Liang Hwu, Pi-Chuan Fan, Ni-Chung Lee, Lee-Jun Wong, Jinglan Zhang, Taosheng Huang

## Abstract

With very few exceptions, mitochondrial DNA (mtDNA) in humans is transmitted exclusively from mothers to their offspring, suggesting the presence of a strong evolutionary pressure favoring the exclusion of paternal mtDNA. We have recently shown strong evidence of paternal mtDNA transmission. In these rare situations, males exhibiting biparental mtDNA appear to be limited to transmitting just one of the mtDNA species to their offspring, while females possessing biparental mtDNA populations consistently transmit both populations to their offspring at a very similar heteroplasmy level. The precise biological and genetic factors underlying this unusual transmission event remain unclear. Here, we have examined heteroplasmy levels in various tissues among individuals with biparental inheritance. Our results indicate that individuals with biparental mtDNA have remarkable inter-tissue variability in heteroplasmy level. At the single-cell level, paternal mtDNA heteroplasmy in sperm varies dramatically, and many sperm possess only one of the two mtDNA populations originally in question. These results show a fundamental, parent-of-origin difference in how mtDNA molecules transmit and propagate. This helps explain how a single population of mtDNAs are transmitted from a father possessing two populations of mtDNA molecules, suggesting that some mtDNA populations may be favored over others when transmitted from the father.

## INTRODUCTION

Mitochondria are organelles which play a critical role in the functioning of the cell, both by providing energy in the form of ATP and by producing a variety of biosynthetic products required for normal development and cellular functions. Mitochondrial dysfunction is increasingly thought to be a major cause of age-related illnesses and progressive neurodegenerative disorders such as Parkinson’s and Alzheimer’s diseases [1]. While most of the genes required for the function of a mitochondrion are encoded by the nuclear genome, a small number of proteins, 22 transfer RNAs (tRNAs), and 2 ribosomal RNAs (rRNAs) are encoded in the small (∼16.5 kb) circular mitochondrial genome that resides within the mitochondrion itself. Mitochondria and the mitochondrial genome (also known as “mtDNA”) appear to have been derived from an ancestral bacterial symbiote. The mtDNA has been shown to be critical for mitochondrial energy production [2]; consequently, mutations in any of its genes can potentially have severe and life-threatening consequences [3].

In humans and most other multicellular organisms, only maternal mtDNA is present in the offspring [4, 5], and while a small amount of paternal mitochondria are initially transferred to the egg, they are actively eliminated via a variety of mechanisms [6–13]. As a result of this paternal exclusion [14], the mtDNA molecules carried by matrilineal individuals are almost identical, and most likely all nucleotide positions are in a state of “homoplasmy”. However, *de novo* mutations can occur in maternal germlines and during somatic development [15]. For these individuals, some of their tissues may contain a mixture of two or more different mtDNA molecules, a state commonly referred to as “heteroplasmy” [16]. While some of these mtDNA variants are apparently benign, others may lead to mitochondrial dysfunction and disease. The severity of disease presentation generally correlates with the identity of the variant and its level of heteroplasmy [17]. For example, due to the random skewing of mtDNA during oocyte development, an overtly healthy mother with a low frequency of deleterious mtDNA can pass on much higher levels of a particular variant to her offspring, thereby increasing the risk of disease in her offspring [18]. Moreover, the levels of heteroplasmy can be variable among different tissues due to random segregation, cell differentiation and tissue-specific energy demand, creating the potential for mitochondrial dysfunction that is isolated to particular tissues or organs [19, 20]. Lastly, there is some evidence that even heteroplasmy for benign variants can be associated with a phenotype [21]. In *C. elegans*, persistence of paternal mitochondria results in a 23-fold increase in embryonic lethality [8]. However, the health of individuals conceived via ooplasm donation [22, 23] appears to contradict inherent pathogenicity for heteroplasmy. All of these factors potentially contribute to the severity of mitochondrial disease, thereby creating unique challenges for better understanding, diagnoses and treatments of mitochondrial diseases.

Recently, we have discovered several rare, familial cases of biparental mtDNA transmission in humans [24]. These rare biparental mtDNA transmission cases provide valuable opportunities to explore the bases of the central dogma of mitochondrial genetics. In this study, we examined the dynamics of heteroplasmy in previously identified biparental mtDNA transmission families. In particular, we have focused our attention on a male in one family who, despite possessing two populations of mtDNA, was observed to transmit only one of these populations to his child. By undertaking a careful examination of heteroplasmy levels in various tissues from this individual and his child, we have sought to obtain a better understanding of the biological and genetic factors that have allowed for the extraordinary transmission of an mtDNA sequence from a father to his child. In addition, we describe three new families which provide additional evidence of biparental mtDNA inheritance and reveal new insights into the nature of this process. One of these families has a very unusual mtDNA deletion and also showed the transmission of only one of the two paternal mtDNA populations. Together, these new results demonstrate that there is a striking degree of cell-to-cell and tissue-to-tissue variability in heteroplasmy levels in individuals with paternally inherited mtDNA populations. Heteroplasmies transmitted through the paternal germline appear to be highly unstable and may be subject to selection favoring some mtDNA populations over others, while heteroplasmies transmitted through the maternal line are highly consistent and stable. Together, the data from these families demonstrate that even under conditions that permit biparental inheritance, there is a fundamental difference in the mechanisms governing the transmission of mtDNA in the maternal and paternal germlines, as well as between the germline and somatic cells.

## RESULTS

### Variations in heteroplasmy levels observed among tissues collected from individuals with biparentally inherited mtDNA

We recently reported multiple unrelated families with paternally transmitted mtDNA [24]. In this report, we noticed two male individuals (II-4 in from Family A and II-3 from Family B) who possessed two populations of variant mtDNA from a prior biparental transmission event, yet only passed one of those two populations on to their offspring. This raised a number of questions as to how one mtDNA haplotype can be excluded by the father while still transmitting the other haplotype, especially in light of the fact that females in these families (and all six families examined so far) who carry mixed populations of mtDNA transmit both populations to their offspring. In light of these observations, we decided to undertake a closer examination of selected members of Family A.

II-4 in Family A carries two populations of mtDNA. However, only one population of mtDNA was passed to his child, III-6. To examine tissue-related differences in heteroplasmy in individuals (II-3, II-4 and III-6) with biparentally inherited mtDNA, a variety of tissue samples were obtained from these individuals. These samples were sequenced as previously described [24–28]. The results obtained were compared to identify changes in heteroplasmy frequency for maternally versus paternally derived mtDNA and their haplogroups.

For II-3 in Family A, their percentage of paternal mtDNA in blood was approximately 36%. In contrast, paternal mtDNA was present at a frequency of less than 2% in their saliva sample (**Fig. S1**). For II-4 from Family A, the percentage of paternal mtDNA in blood was similar, with approximately 40% heteroplasmy detected by PCR-NGS (**Fig. 1B**). In the other tissues tested from II-4, paternal heteroplasmy was variable from 0% in urine, 13% in fibroblasts, and 20% in sperm. This pattern of lower paternal heteroplasmy was also observed in II-4’s child, III-6, who carries 40% paternal heteroplasmy in blood, yet less than 2% heteroplasmy in saliva and urine (**Fig. S2**). It should be noted that no paternal heteroplasmy exceeded 40%, thereby demonstrating that (at least so far) dominance of the maternal haplogroup was maintained even under conditions of biparental transmission.

**Figure 1.**
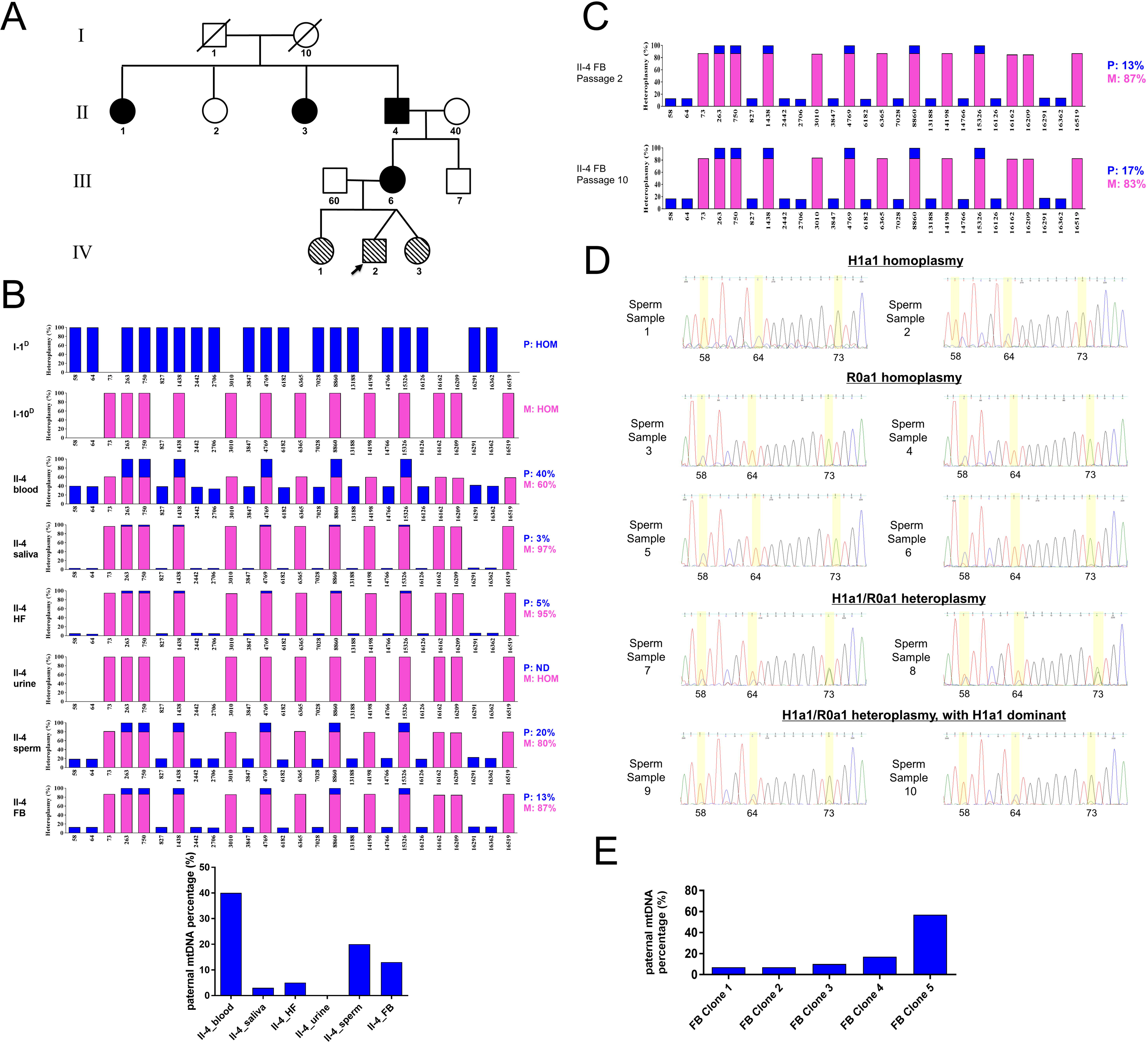
Tissue-related variation in heteroplasmy levels for maternal (H1a1) and paternal (R0a1) haplogroups in Patient II-4 of Family A. (**A**) Pedigree of Family A. The black-filled symbols indicate the four family members (II-1, II-3, II-4, and III-6) that show biparental mtDNA transmission. The diagonal-filled symbols indicate the three family members (IV-1, IV-2, and IV-3) who carry a high number and level of mtDNA heteroplasmies which underwent normal maternal transmission. (**B**) Variable heteroplasmy levels detected by PCR-NGS in blood, saliva, hair follicle (HF), urine, sperm, and fibroblast (FB) samples collected from Patient II-4. (**C**) Heteroplasmy drift over time increases the proportion of the paternal haplogroup in primary fibroblast cells derived from Patient II-4. (**D, E**) Single cell-derived DNA samples from Patient II-4 were sequenced to measure cell-to-cell variability in heteroplasmy levels. Sanger sequencing for single-sperm samples (**D**) and PCR-NGS for colonies derived from individual primary fibroblast cells (**E**) show marked differences in heteroplasmy levels, even for cells derived from the same tissue.

### Heteroplasmy drift in extended fibroblast cultures

Our tissue-specific sequencing data confirmed that mtDNA heteroplasmy levels can dramatically diverge between different tissues in the same individual. This phenomenon has generally been attributed to the assumption that mitochondria (and their associated mtDNAs) are randomly segregated during cell division. As a result, mtDNA molecules are differentially distributed among different precursor cells during development. This creates a genetic drift effect whereby the frequency of mtDNA changes with proliferation over time. Rapid shifts in heteroplasmy have been observed in cell culture [29], as well as in the children of mothers carrying disease-causing mtDNA variants [30]. However, whether such changes are truly random, or whether they involve a selection favoring one mtDNA molecule over another, remains unknown in the majority of cases. The co-existence of two widely divergent mtDNA molecules generated by biparental transmission creates a unique opportunity to examine this question in a novel context. Therefore, directional changes in paternal heteroplasmy were examined in primary dermal fibroblasts obtained from II-4 of Family A over time. A slight, yet consistent, increase in the frequency of paternal mtDNA was observed over eight passages. For example, the initial paternal mtDNA frequency was ∼12% at Passage 2, and it increased to ∼17% at Passage 10 (**Fig. 1C**). While this result is consistent with a preferential increase in the paternal haplogroup (R0a1) over time, the magnitude of the effect is quite small, and additional experiments and samples will be required to confirm and further generalize this result.

### Heteroplasmy exhibits cell-to-cell mosaicism in sperm and fibroblast cells

Tissue-specific sequencing can provide valuable information regarding heteroplasmy dynamics within a particular individual. However, the data only represent a general summary of the events occurring at the cellular level. In theory, there can be a wide variety of heteroplasmies present within a tissue, even across consecutive tissue samples. Thus, determining the exact level of cell-to-cell heteroplasmic variability in different tissues may provide valuable insight into the mechanisms guiding mtDNA transmission, particularly for biparental mtDNA inheritance.

Ten single-sperm samples from II-4 in Family A were therefore subjected to Sanger sequencing. The overall heteroplasmy level for the paternal haplogroup in these sperm samples was approximately 20% (**Fig. 1B**), and a wide range of heteroplasmic variants were clearly observed over these single-sperm samples (**Fig. 1D**). Specifically, homoplasmic or nearly homoplasmic variants for R0a1 (the paternal haplogroup) were identified (n = 4), homoplasmic or nearly homoplasmic variants for H1a1 (the maternal haplogroup) were identified (n = 2), and heteroplasmic variants were also identified (n = 4). Two of the heteroplasmic samples were biased heavily toward H1a1, suggesting an even split between R0a1 and H1a1 in these ten sperm samples. While this ratio differs from the overall paternal heteroplasmy level of 20%, this difference may be due to the relatively small sample size of sperm samples that were analyzed and the presence of other cells in the total sperm sample (e.g. leukocytes and immature germ cells), which could potentially exert an outsized effect on the bulk samples due to the low mtDNA copy number found in mature sperm cells. By contrast, leukocyte-derived mtDNA was unambiguously excluded from our single sperm samples, as we sorted individual sperm cells using a piezo micromanipulator and microscope. Regardless, it is clear that a sizeable percentage of the germ cells examined are effectively homoplasmic, and both maternal and paternal haplogroups are represented.

Next, we examined a somatic cell type to compare it with the single-sperm samples. Briefly, clonally derived colonies were generated from primary dermal fibroblasts obtained from II-4 from Family A. Similar to the sperm sample results from the same individual, NGS sequencing showed paternal heteroplasmy levels for some fibroblast colonies to be well outside the range predicted, with one colony showing paternal heteroplasmy as high as 57% (**Fig. 1E**). Meanwhile, the paternal heteroplasmy levels for other colonies from II-4 were virtually unchanged from the 13% heteroplasmy level identified with sequencing of the bulk fibroblast samples (**Fig. 1B**). While these data do not represent a comprehensive analysis of all cell types, the results obtained do derive from widely divergent cell types, and they confirm that mtDNA mosaicism is likely to be a general phenomenon throughout the body.

### New evidence of biparental mtDNA inheritance

Recently, we have discovered three additional families (Family D, Family E and Family F) with biparental mtDNA inheritance. These cases provide further evidence regarding biparental mtDNA transmission in humans, and they also exhibit some interesting difference from our previously observed cases. Thus, new insights into the unique process of biparental mtDNA inheritance were obtained.

### Family D: A long-lasting transmission of biparental mixture in a normal, maternally inherited manner

Proband IV-7 (age: 16 years and 5 months) of Family D (**Fig. 2A**) received a diagnosis of white matter disease involving the spinal cord and cerebellum, progressive neurologic illness, and gait abnormality secondary to cerebellar lesions. She was previously evaluated for visual disorders such as hypermetric saccades and end-gaze nystagmus (consistent with cerebellar disease), as well as temporal optic disc pallor in both eyes. Regarding the latter, spectral domain-optical coherence tomography (SD-OCT) showed the retinal nerve fiber layer to be stable and unchanged. Other diagnoses for this patient included: allergic rhinitis, oropharyngeal dysphagia, abdominal pain, emesis, chills, headaches, tiredness, and weakness.

**Figure 2.**
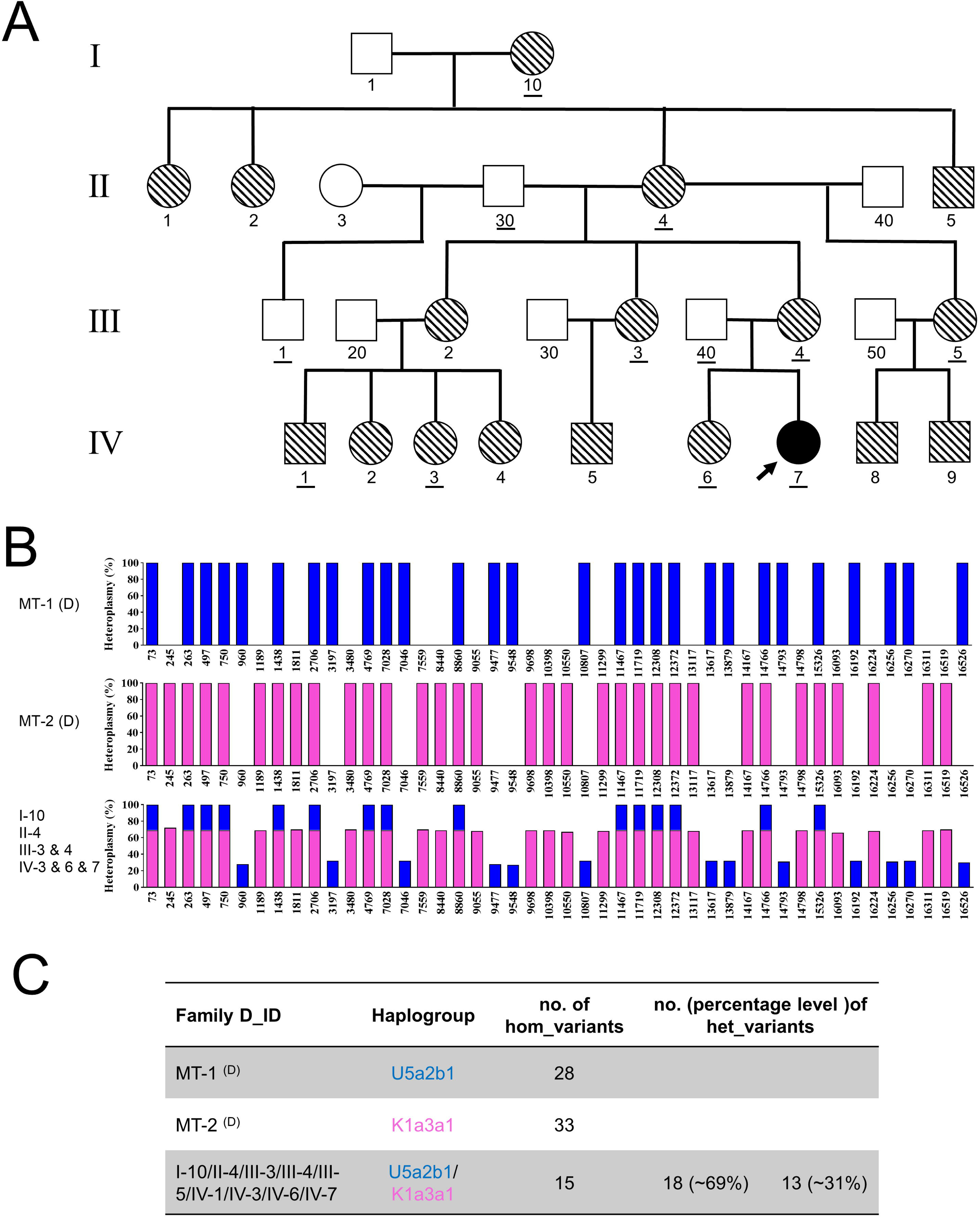
Cross-generation stability of a likely biparental mtDNA transmission event in Family D. (**A**) Pedigree of Family D. The solid-filled symbol indicates the proband (IV-7), and the diagonal-filled symbols indicate the other seventeen family members (I-10, II-1, II-2, II-4, II-5, III-2, III-3, III-4, III-5, IV-1, IV-2, IV-3, IV-4, IV-5, IV-6, IV-8 and IV-9) who carry a high number and level of mtDNA heteroplasmies which underwent normal maternal transmission. No paternal transmission event was directly observed in this family. The IDs of the twelve family members who underwent whole mtDNA sequencing are underlined; all other genotypes are inferred. (**B**) Schematic diagram of the mtDNA genotypes defined by the homoplasmic and/or heteroplasmic variants identified from a comparison with a reference mitochondrial genome. Since the paternal or maternal origin of the two haplogroups (MT-1 and MT-2) cannot be unambiguously established from the current data, blue and pink bars are used to represent the two distinct haplogroups detected in the heteroplasmic individuals. Stacked blue and pink bars indicate variants that are homoplasmic because they are shared by both haplogroups. Entries labeled (D) represent the deduced mtDNA genotypes of I-10’s parents. The heteroplasmy levels determined from the II-4 sample are shown as a representation of the U5a2b1/K1a3a1 heteroplasmy being transmitted in this family. (**C**) Summary of the haplogroup and mtDNA variant numbers in Family D.

The patient developed dysarthria recently and difficulty walking for any distance without the use of a wheelchair. The patient was previously diagnosed as exhibiting a delay in development, although her current behavior and social skills are typical for her age. She also appears to have no major cognitive issues. The patient has further experienced an increase in spasticity and loss in range of motion for her ankles and knees. Following immunotherapy, the patient exhibited some improvement, although the therapy was discontinued due to concern for further decline. In previous MRI scans, abnormalities within the spinal cord were observed which were consistent with demyelinating disease. However, subsequent MRI exams were stable, with no new areas of involvement identified.

The patient’s condition has defied a genetic diagnosis despite application of broad genetic testing. Moreover, a Trio analysis of exome sequencing data revealed no clinically significant results. In addition, panels for Friedreich ataxia, spinocerebellar ataxia, and other forms of ataxia produced negative results. Thus, patient IV-7 was referred to CCHMC to be evaluated for a potential mitochondrial disease.

Mitochondrial sequencing of proband IV-7 and alignment of these data to the human mitochondrial sequence reference, NC_012920.1, revealed 15 homoplasmic variants (indicated with pink or blue bars; **Fig. 2B**) and 31 heteroplasmic variants, none of which are predicted to be pathogenic variants. Among the former, eleven were common to both maternal (K1a) and paternal (U5a) haplogroups. Among the latter, 18 variants exhibited ∼60% heteroplasmy and were associated with the K1a3a1 haplogroup (pink bar), while the other 13 variants exhibited approximately 40% heteroplasmy and were associated with the U5a2b1 haplogroup (blue bar).

Based on our previous experience with the presence of two distinct haplogroups in patient mtDNA, we initially predicted that biparental transmission occurred in proband IV-7. Moreover, the haplogroups of IV-7 appear to derive from the father and from the mother. However, sequencing of the father (III-40) revealed that he did not contribute to either haplogroup. Instead, it was confirmed that the mother (III-4) possessed the same heteroplasmies at comparable levels to both the proband and the proband’s sister (IV-6), thereby implying that the putative biparental transmission event likely occurred in an earlier generation.

Consequently, additional family members from various generations (I-10, II-4, III-3, III-4, III-5, IV-1 and IV-3) were recruited to undergo sequencing and identify the original biparental transmission event(s). However, no such event was identified. Instead, the mixture of U5a2b1/K1a3a1 haplogroups was found to originate at least as far back as the proband’s great-grandmother (I-10), and it has been passed on at more or less the same heteroplasmy levels (anywhere from ∼70% to 45% K1a3a1 heteroplasmy and ∼30 to 55% U5a2b1 heteroplasmy) in all subsequent maternal descendants. The consistent appearance of this specific admixture of haplogroups strongly implies that a paternal transmission event occurred in an ancestor of the proband’s great-grandmother (I-10), as this would appear to be the most likely explanation for how two distinct haplogroups could be made to cosegregate in such a manner. This presumptive biparental inheritance event was then transmitted through subsequent generations in a normal, maternally inherited manner. Thus, it appears that the products of paternal transmission can persist through many generations (presumably ending when one of the two haplogroups is randomly lost due to a meiotic bottleneck), thereby having a long-lasting genetic effect within a family or population.

### Family E: Paternal transmission of mtDNA carrying a large deletion

Family E (**Fig. 3A**) is particularly interesting, as it involves the paternal transmission of an mtDNA molecule carrying a large deletion. Proband (III-1) is an 18-year-old female who manifested several pathologies potentially related to mitochondrial dysfunction. These pathologies included: optic nerve atrophy, bilateral basal ganglion calcification, migraines, vertigo, muscle weakness, inflammatory bowel disease, hypothyroidism, congenital adrenal hyperplasia, arthritis, and vasculitis. In addition, the proband’s father (II-1) and brother (III-2) were previously diagnosed with migraines and high myopia. The father (II-1) also experienced frequent diarrhea. Several members of the proband’s extended family presented with diabetes, including two maternal uncles (II-2 and II-4), an aunt (II-3), and the grandmother (I-10). I-10 also suffered from retinopathy (presumably as a complication from diabetes), while I-10 and I-1 presented with hypertension. Thus, III-1 was referred to CCHMC for evaluation.

**Figure 3.**
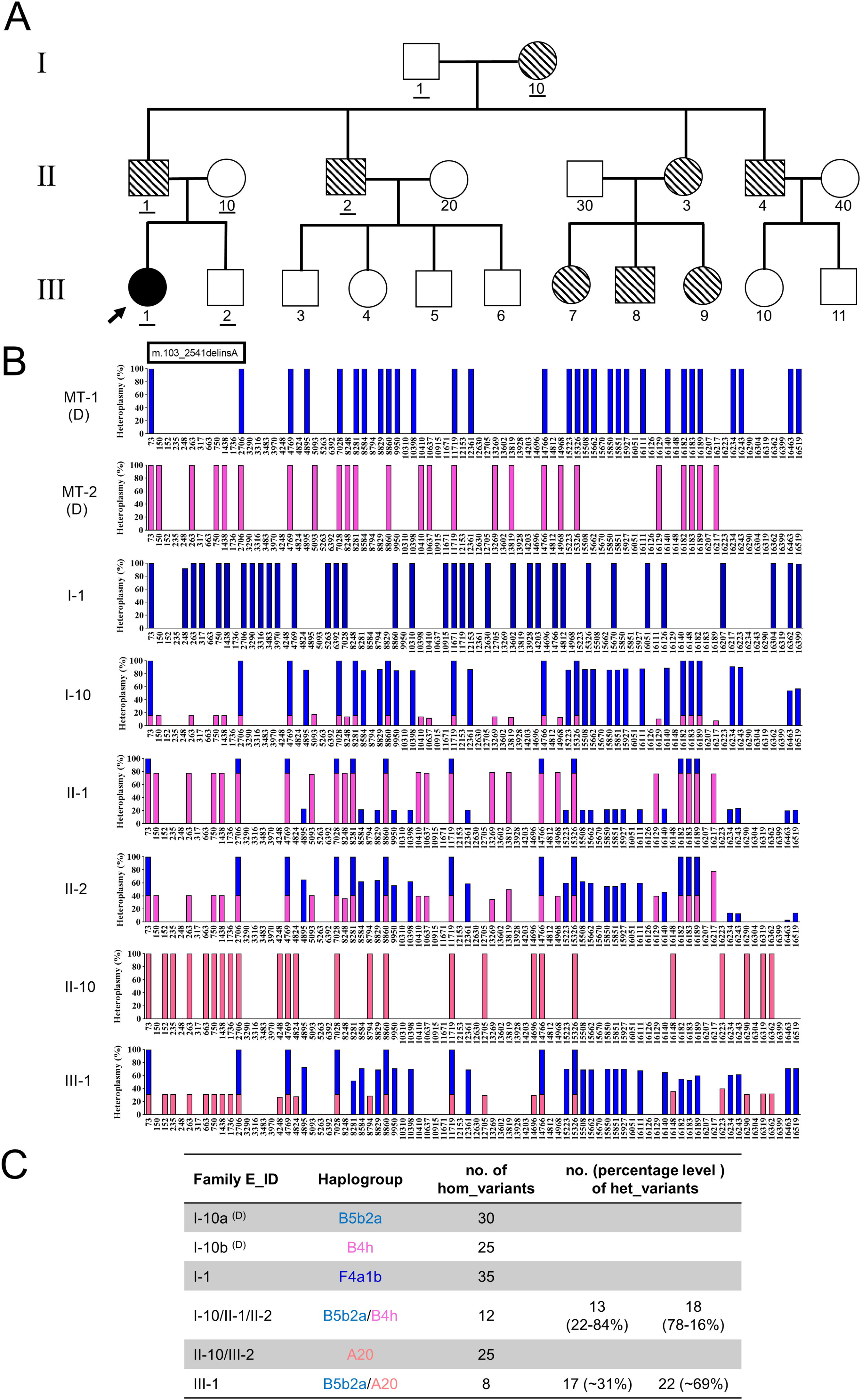
Paternal transmission of a large deletion allele in Family E. (**A**) Pedigree of Family E. The black-filled symbol indicates the family member (III-1) that exhibits biparental mtDNA transmission. The diagonal-filled symbols indicate the eight family members (I-10, II-1, II-2, II-3, II-4, III-7, III-8, and III-9) who carry a high number and level of mtDNA heteroplasmies which underwent normal maternal transmission. The IDs of the twelve family members who underwent whole mtDNA sequencing are underlined; all other genotypes are inferred. (**B**) Schematic diagram of the mtDNA genotypes defined by the homoplasmic and/or heteroplasmic variants identified from a comparison with a reference mitochondrial genome. Blue bars represent the genotype of paternally derived mtDNA (i.e. haplogroup B5b2a), while pink and orange-red bars represent maternally derived mtDNA. Since the paternal or maternal origin of the two haplogroups (MT-1 and MT-2) in I-10 cannot be unambiguously established from the current data, blue and pink bars are used to represent her two distinct haplogroups. The dark blue bars represent the homoplasmic genotype of the grandfather, I-1. The entries labeled “D” represent the deduced mtDNA genotypes of I-10’s parents. (**C**) Summary of the haplogroup and mtDNA variant numbers in Family E.

Whole exome sequencing of the parent-proband trio was performed and variants in the *CYP21A2* gene were identified. These variants included compound heterozygous pathogenic variants, c.518T>A (p.I173N) and c.955C>T (p.Q319X), and these were confirmed with Sanger sequencing. Sanger sequencing also showed that the mother (II-10) is a heterozygous carrier of the c.518T>A (p.I173N) allele, while the father (II-1) is heterozygous for the c.955C>T (p.Q319X) allele. Mutations in *CYP21A2* can cause adrenal hyperplasia 3 [MIM:201910], a common recessive disease that is due to defective synthesis of cortisol. Congenital adrenal hyperplasia is characterized by excess androgen that leads to ambiguous genitalia in affected females, while rapid somatic growth during childhood with premature closure of the epiphyses and short adult stature are observed in both affected sexes. A heterozygous c.2817dupT (p.H940fs) variant in the *NBAS* gene, which is likely pathogenic, was also detected. Defects in NBAS cause short stature, optic nerve atrophy, and Pelger-Huet anomaly (SOPH) [MIM:614800]. The latter is an autosomal recessive syndrome characterized by severe postnatal growth retardation, facial dysmorphism with senile face, small hands and feet, normal intelligence, abnormal nuclear shape in neutrophil granulocytes (Pelger-Huet anomaly), and optic atrophy with loss of visual acuity and color vision.

Since whole exome sequencing failed to provide a complete explanation of the proband’s symptoms, whole mtDNA sequencing was subsequently performed for the proband-parent trio. Alignment of whole mtDNA data to the human mitochondrial reference sequence, NC_012920.1, identified 8 homoplasmic variants (orange-red plus blue bars, **Fig. 3B**) and 39 heteroplasmic variants. The 8 homoplasmic variants are predicted to have been inherited from the proband’s parents. For the heteroplasmic variants, a segregation analysis showed that 17 have a 31% heteroplasmy level (labeled with orange-red bar) and were maternally inherited. Meanwhile, the remaining 22 heteroplasmic variants had a 69% heteroplasmy level and were inherited from the proband’s father (blue bar). The proband’s father was also found to carry multiple heteroplasmic mtDNA variants similar to the proband.

Other family members were recruited to further delineate the mtDNA transmission events that had occurred in previous generations. The results indicate that the heteroplasmic condition of II-1 was transmitted to him from his mother, who was also heteroplasmic for B5b2a (blue bars) and B4h (pink bars). On the other hand, II-1 received no mtDNA contribution from his father (I-1), who is homoplasmic for the F4a1b haplogroup (dark blue bars, **Fig. 3B**). Sequencing showed that a paternal uncle of the proband (II-2) also carried both the B5b2a and B4h haplogroups, confirming that I-10 is the source of this heteroplasmy.

In addition to the previously described 22 heteroplasmic variants, NGS showed that both the proband and her father carry the same large deletion variant in their mitochondrial genome, m.103_2541delinsA, at low heteroplasmy. This deletion, which is predicted to be a pathogenic variant, was also detected in the grandmother’s (I-10) DNA by mtDNA sequencing, as well as in the DNA of one of II-1’s siblings (II-2) (**Fig. 3A**). Given the extremely low probability that this exact deletion would appear independently in all of these individuals, these results indicate that this deletion was passed on as part of the B5b2a haplogroup from the grandmother (I-1) to the proband’s father (II-1), and then passed on via paternal transmission from II-1 to III-1. Thus, this mtDNA deletion provides the strongest single piece of genetic evidence to date regarding the ability of a human male to transmit an mtDNA sequence variant to one of his offspring.

### Family F: Two more likely cases of father to offspring mtDNA transmission

The proband (II-1) of Family F is a 30-year-old female who presented to the Mitochondrial Genetics Clinic at CCHMC with clinical phenotypes suggestive of mitochondrial disease. The patient was apparently healthy until around the age of 25 years, when she developed a *C. difficile* infection and started becoming fatigued easily. Other symptoms exhibited by the proband include cardiomyopathy, postural orthostatic tachycardia syndrome, muscle weakness, migraines, mastocytosis associated with GI symptoms, iron and B12 deficiencies, low phosphorus levels, chronic hives and shortness of breath with exertion. In addition, other family members have been diagnosed with “mitochondrial diseases.” Specifically, the proband’s mother has also shown signs of chronic fatigue, and her five-year-old daughter has shown signs of delayed growth.

As the first step of evaluation, the proband’s lactate and pyruvate levels, as well as the ratio of lactate to pyruvate, were all apparently normal. However, her mitochondrial whole genome sequencing showed that she has multiple heteroplasmic variants, none of which are known to be pathogenic. Since this is a pattern that we have observed before with biparental mtDNA inheritance, additional family members were enrolled in the study to determine the transmission pattern of mtDNA haplotypes within her immediate family. Sequencing of the proband’s three children (III-1, III-2 and III-3), her brother (II-2), and her mother (I-1) suggests that biparental mtDNA transmission has indeed occurred in this family (**Fig. 4A**). Specifically, our analysis identified 9 homoplasmic variants (pink plus blue bars, **Fig. 4B**) and 13 heteroplasmic variants in the mtDNA of the proband, as well as her brother (II-2) and all three of her children (III-1, III-2 and III-3). The 9 homoplasmic variants are predicted to have been inherited from both the mother and the father of the proband. However, for the heteroplasmic variants, a segregation analysis showed that 3 variants have an average heteroplasmy around 81% (labeled with pink bars, **Fig. 4B**) and were inherited from the mother (I-1), who is homoplasmic for the H7b haplogroup. Meanwhile, the remaining 10 heteroplasmic variants had ∼19% heteroplasmy level (blue bars, **Fig. 4B**) and were not present at all in the mother; these results suggest that the proband has inherited mtDNA from her father (I-10), who is deduced to have possessed the U haplogroup. However, I-10 is not available for confirmation. Overall, this analysis indicates that the proband and her brother both inherited the U haplotype from their father at approximately the same level (19% for II-1 and 20% for II-2), resulting in a heteroplasmic mixture of the U and H7b haplogroups. This suggests that paternal transmission events have occurred in this family. Furthermore, we can see that the proband passed on this mixture to all three of her children, again at a relatively consistent level (12% to 19% heteroplasmy for haplogroup U). This demonstrates once again that when a paternal transmission results in a female offspring with mixed haplotypes, that she will invariably pass on that mixture to all of her offspring.

**Figure 4.**
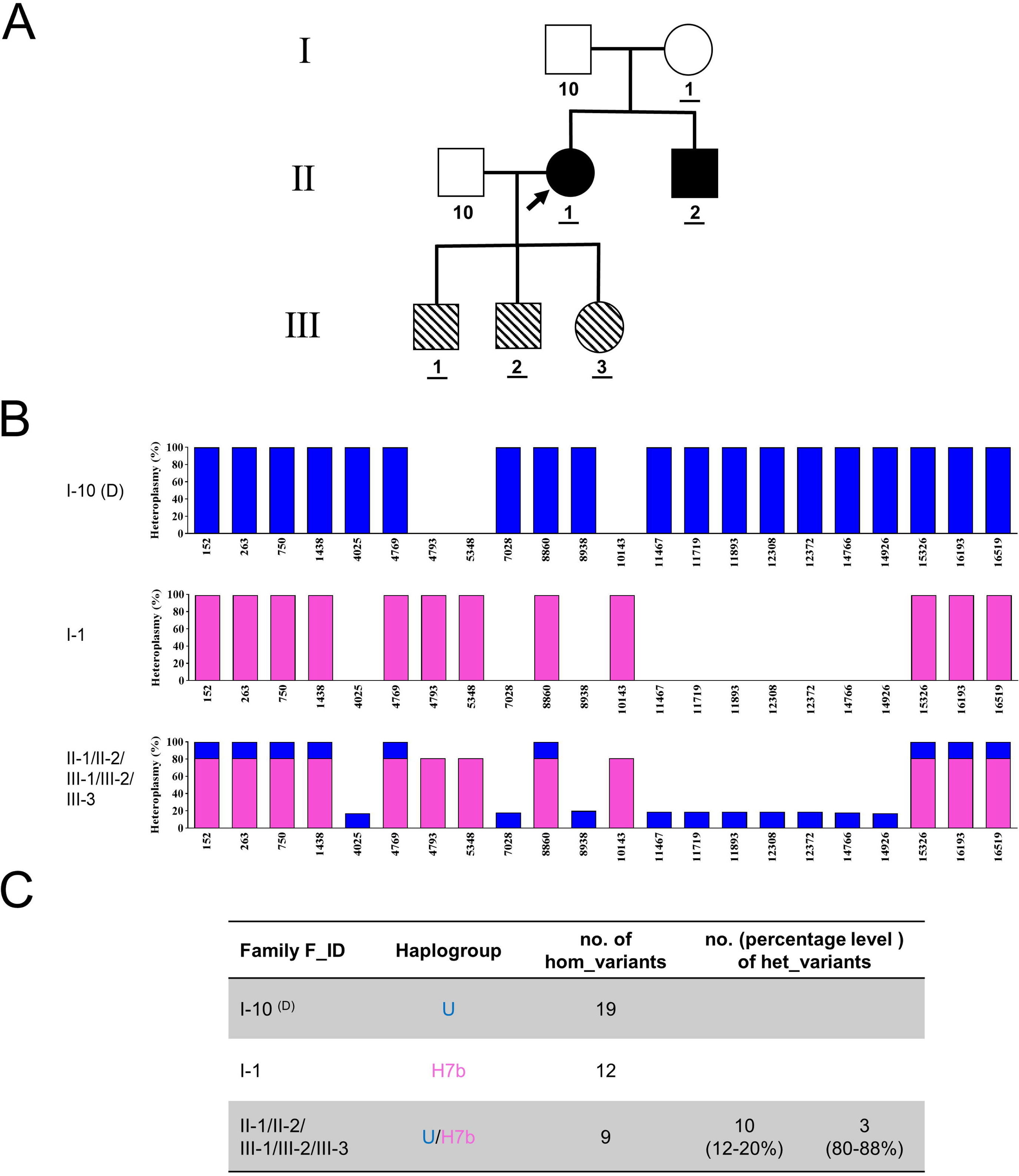
Two paternal transmission events observed in Family F. (**A**) Pedigree of Family F. The black-filled symbols indicate the family members (II-1 and II-2) that are deduced to have inherited the U haplogroup from their father (I-10), as their mother (I-1) is homoplasmic for the H7b haplogroup. The diagonal-filled symbols indicate the three family members (III-1, III-2, and III-2) who carry a high number and level of mtDNA heteroplasmies which underwent normal maternal transmission. The IDs of the twelve family members who underwent whole mtDNA sequencing are underlined; all other genotypes are inferred. (**B**) Schematic diagram of the mtDNA genotypes defined by the homoplasmic and/or heteroplasmic variants identified from a comparison with a reference mitochondrial genome. Blue bars represent the genotype of the paternally derived mtDNA (haplogroup U), while pink bars represent the maternally derived mtDNA (haplogroup H7b). Stacked blue and pink bars indicate variants that are homoplasmic because they are shared by both the paternal and maternal haplogroups. The entry labeled “D” represents the deduced mtDNA genotype of the grandfather (I-10). (**C**) Summary of the haplogroup and mtDNA variant numbers in Family E.

## DISCUSSION

The results of this study demonstrate that heteroplasmies inherited as a result of biparental inheritance exhibit different modes of transmission in subsequence generations based on the parental origins. We observed that these heteroplasmies show relatively consistent transmission of both mtDNA populations in the female germline across generations. On the other hand, paternal transmission appears to favor exclusive transmission of one mtDNA population over the other when the father is heteroplasmic. Heteroplasmy levels were also observed to differ wildly between cells residing within the same tissue, consistent with replicative segregation, one of the characteristics of mitochondrial genetics. While the precise molecular mechanisms mediating biparental mtDNA transmission events remain to be fully elucidated, several important insights can be inferred from the present results.

First, the high degree of variability in the mtDNA genotypes of the individual sperm cells is truly fascinating and may explain a great deal about the transmission pattern observed in the paternal line. A similar situation has been previously described in mice, where significant variance in heteroplasmy was recently confirmed at the single-cell level in both oocytes and somatic cells [31]. This genotypic variability appears to be the result of the paternal “mtDNA bottleneck” effect, similar to the well-known maternal mtDNA bottleneck which contributes to the dramatic shifts in mtDNA heteroplasmy that if often observed from mother to offspring. This effect was traditionally ascribed to a massive reduction in mtDNA copy number that occurs during early development [32], with the implication that this copy number restriction leads to genetic drift and changes in mtDNA variant frequency. However, a contradictory view has been proposed whereby the bottleneck effect is largely independent of mtDNA copy number reduction, and instead, is primarily due to random partitioning of mtDNA during cell division [33, 34]. The results of the present study favor heteroplasmy combined with random partition (since a range of possible heteroplasmies were observed within a small sample of ten individual sperm cells), but there is a possibility of selective forces that drive the sperm toward homoplasmy. Even more intriguing is the possibility that sperm possessing a specific mtDNA may have an advantage in the fertilization. It is also striking that the sequencing of single fibroblast clones revealed a similar dynamic in a somatic tissue, with distinct variations in paternal heteroplasmy levels observed from one cell to the next. Previous studies have described cell-to-cell heteroplasmic stability in cell lines carrying more usual forms of heteroplasmy (i.e., *de novo* mutations and maternally inherited mtDNA variants) (24). This stability has been attributed to intercellular exchanges of mtDNA molecules [35]. However, the rigorous investigation undertaken here clearly indicates that mosaicism in heteroplasmy levels is likely to be a common occurrence, thereby providing a plausible mechanism for the shifts in mtDNA heteroplasmy observed in both our patient samples and with extensive *in vitro* cell culturing.

Another interesting implication of the variable heteroplasmy observed in the sperm samples examined is apparent from the pedigree of Family A. As shown in our previous publication, II-4 only transmitted mtDNA to one of his offspring (III-6), while the other offspring received maternal mtDNA in the usual manner (see **Fig. 1A, B**, and [24]). The presence of paternal mitochondria in II-6, but not in III-7 seem to imply that some sperm carry the paternal transmission trait, while others do not. Assuming the paternal transmission trait is caused by mutation of a nuclear locus involved in tagging or signaling paternal mitochondria for destruction, this behavior could be explained by the haploid nature of the sperm cell, with some sperm carrying the permissive mutant allele while other sperm carry the nonpermissive wildtype allele. Furthermore, of the two haplogroups carried by II-4, only the paternally inherited haplogroup was transmitted to III-6. Based on the results obtained from the single-sperm sequencing analysis, it appears that this transmission pattern is likely the result of fertilization by sperm homoplasmic for the paternally inherited haplogroup. It remains to be determined whether this fertilization pattern was random or the result of biological selection in favor of the R0a1 haplogroup. It is also noteworthy that II-1 in Family E transmitted one particular haplogroup to his daughter, and that the transmitted haplogroup carried a deletion that may confer a size-based proliferative advantage. Both results could imply the involvement of some sort of selective process. At least one group has come out in favor of this model, arguing that if one of the paternal mtDNA possesses a proliferative advantage over the other, that this fact combined with the random partitioning of mtDNA during sperm development would be sufficient to create the autosomal-like dominant transmission pattern we have observed in these families for paternal mtDNA [36]. In support of this hypothesis, at least one study has shown that mice carrying heteroplasmic mixtures of mtDNA from different genetic backgrounds demonstrate preferential segregation of some mtDNAs over others [21]. While we have previously described our own reasoning for why a permissive autosomal locus is still likely to be necessary for paternal mtDNA transmission to occur [37], future work will help settle this question, as well as clarify the relative contribution of random versus selective factors in the phenomenon of biparental mtDNA transmission.

According to the theory that an mtDNA bottleneck exists during early development, heteroplasmies generated through biparental inheritance should be rather unstable, with frequencies shifting quite dramatically across generations. This is apparent in the paternal transmission events, where heteroplasmy shifts quite dramatically across generations. For example, as observed in Family A (**Fig. 1**), only one of the two heteroplasmies observed in II-4 is passed on to his daughter (III-6). However, in contrast to the instability of paternally transmitted heteroplasmies, once a heteroplasmy becomes established in a daughter, it appears to become remarkably stable in composition. In fact, these mixed mtDNA populations appear to pass from mother to child as with any other mtDNA population, as observed in III-6 from Family A (**Fig. 1** and [24]). Moreover, the clearest example of heteroplasmic stability is observed in Family D (**Fig. 2**), where a similar mixture and frequency of haplogroups can be observed in multiple individuals across at least four generations. How this stability observed in Family D is maintained for so long (given the randomness imposed by the mtDNA bottleneck) remains an important question for future investigation, particularly in light of the more dramatic heteroplasmy shifts observed in Family E (**Fig. 3**).

There also remains the question of the ultimate fate of the mixed haplotypes created by biparental mtDNA transmission. It has been shown in the guppy species, *Poecilia reticulata*, that complex heteroplasmies of up to nine different haplotypes have been not only detected, but were shown to be passed from mother to offspring [38]. However, it remains unclear why complex mixtures of haplogroups are not observed more often in humans. As one group has pointed out, if the ability to transmit paternal mtDNA sequences can be carried within a family, this would be expected to result in the linear accumulation of several mtDNA haplotypes over time [39], which is strikingly similar to the situation described above for *Poecilia reticulata*. However, two possible explanations come to mind for why we have not observed this in humans. First, if paternal mtDNA transmission can only occur in situations where a particularly competitive mtDNA is combined with the autosomal dominant “permissive factor,” then this would drastically limit the number of distinct haplotypes that can accumulate over time. Second, it is entirely possible that these mixtures of haplogroups, while stable across a small number of generations, are intrinsically unstable situation in the long term, and that one of the two haplogroups is always lost over evolutionary time due to genetic drift or preferential replication of one of the two haplogroups.

Another fundamental consideration that remains to be resolved is how paternal mtDNA manages to persist long enough to transmit from a father to his daughter, at which point it is then able to follow the maternal inheritance pattern in subsequent generations. As suggested above, it appears likely from the existing family pedigrees that an autosomal locus is required for paternal transmission to occur. Although the identity of that locus remains unclear, it is likely that the gene in question has some involvement in the process by which paternal mitochondria are marked for destruction by mitophagy shortly after fertilization [11, 12, 40]. However, there is a second hurdle that must be crossed by the paternal mtDNA. Given the small number of mtDNA molecules carried by sperm (which about 10,000 times lower than the number carried by the oocyte) [41], paternal mtDNA starts out at an extreme numerical disadvantage relative to the maternal contribution. Even assuming that active mechanisms of paternal mitochondrial elimination (PME) are bypassed in these families, an additional intervention is needed to bring the percentage of paternal mtDNA into the range observed in patients, which is sometimes as high as 40%. One plausible explanation may involve a preferential amplification of paternal mtDNA during early embryonic development. There is already precedence for this situation with maternally inherited mtDNA, where only a subset of mtDNA molecules are allowed to replicate post-fertilization [42]. Furthermore, the reduction from an estimated ∼200,000 copies of mtDNA in the oocyte to the ∼100 to 10,000 copies (“bottleneck effect) typically observed in most somatic cells means that the mitochondria do not normally replicate during a portion of embryogenesis. At the very least, research in mouse embryos has shown that there is normally no mtDNA replication prior to the blastocyst stage [43]. While the mitochondria undergo fission (but not mtDNA replications) ahead of cells division, they remain attached to cytoskeletal elements [44], and this tethering may be associated with the lack of replication. If untethered mitochondria (i.e. paternal mitochondria) are free to divide, then this could explain their overrepresentation in individuals bearing paternal mitochondria. A simple calculation shows that if the number of paternal mitochondria retained in a daughter cell were to double with each cell division, then in 10 cell divisions the paternal to maternal mitochondrial ratio would be 83%, assuming that no maternal mitochondria are free to replicate during the first 10 divisions. This model, while speculative, would be consistent with the results we have observed in these families.

If preferential amplification is enough to explain how paternal mtDNA overcomes its numerical disadvantage, then it’s likely that the identity of the mtDNA itself plays a contributing role in the process, as polymorphic mtDNA sequences (particularly in the D-Loop region) are known to contribute to differences in replication rates between different mtDNAs during mitochondrial replacement therapy [45]. If the same phenomenon holds true in this context, this may be yet another factor contributing to the rarity of paternal mtDNA transmission, as the right confluence of a permissive autosomal locus as well as an mtDNA haplotype capable of overcoming the numerical disadvantage would be required for paternal transmission to be observed. Furthermore, this mechanism may also explain why fathers carrying paternally inherited mtDNA preferentially transmit one of those mtDNAs, as one of the two mtDNA haplotypes may simply outcompete the other haplotype at a key point during germline or embryonic development.

One other important question can be addressed based on the transmission pattern of these mixed populations of biparentally inherited mtDNA. In response to our previous report of paternally transmitted mtDNA sequences [24], it has been proposed that these mitochondrial sequences are not true mtDNA molecules in their normal state, but are instead the result of NUMTs (“nuclear mitochondrial DNA segments”), which are fragments of the mitochondrial genome present as insertions into the nuclear genome [46]. If a large enough concatemer of full-length mtDNA sequences were to insert into the nuclear genome, it might be able to produce a strong enough signal to compete with the hundreds or thousands of copies of genuine mtDNA normally present in the cell. This contention was also raised by a separate letter published in *Frontiers in Genetics* [47], whose authors coined the term “Mega-NUMT” to described such an unusually large NUMT, which would be hundreds of kilobases (if not several megabases) in size. The authors of the original letter proposing this hypothesis have recently presented a poster abstract at the ASHG 2019 meeting that further advances this point [48]. The authors describe a new family, unrelated to any of the ones we have studied, that contains healthy individuals who are also transmitting a mixture of two distinct mtDNA haplogroups. However, they were able to further extend their analysis by creating mtDNA-depleted “ρ0 cells” from one of their patient fibroblast samples, which showed only the presence of a single haplogroup after mtDNA depletion. This suggests that the remaining haplogroup may be associated with a Mega-NUMT, which they estimate to contain anywhere from 45-56 mtDNA copies by ddPCR (implying an insertion of roughly 762 kb to 928 kb). This could represent a novel and unique mechanism, and merits further study going forward. Moreover, as the members of this family are described as “healthy individuals,” this particular finding of biparental mtDNA inheritance would seem to be purely incidental. This raises the possibility that biparental mtDNA inheritance could be more common than previously anticipated, since many such individuals are asymptomatic.

We have offered our own counterpoints and perspectives on the Mega-NUMT hypothesis in a previous letter [49], and the additional families uncovered in the current manuscript allow us to further expand upon this discussion. First, if a Mega-NUMT containing 50+ copies of the full-length mtDNA sequence is indeed possible, the percentage of the Mega-NUMT would vary from one tissue to another, due to differences in mtDNA copy number intrinsic to different types of cells and tissues [50]. This would cause the apparent level of the Mega-NUMT to be higher in tissues where the mtDNA to nuclear DNA ratio is low (e.g. blood) and lower in tissues where the mtDNA to nuclear DNA ratio is higher (e.g. hair). However, this is not what we see in our Family A. For example, the observed heteroplasmy levels in the saliva and hair of II-4 from Family A are almost identical (3% and 5%, respectively), despite the widely divergent mtDNA:nDNA ratios of those two tissues (**Fig. 1B**).

In addition, we continue to observe a pattern across these mixed haplotype families that cannot be easily explained by the Mega-NUMT hypothesis: namely, the maternal transmission of these biparentally generated heteroplasmies in subsequent generations. After all, a NUMT is, by definition, a locus in the nuclear genome, and would be expected to be present as a heterozygous allele in the individuals carrying these apparent heteroplasmies (or in a hemizygous state in the case of a male’s X chromosome). Given those conditions, both males and females would be expected to pass on such an allele approximately 50% of the time if it was located on an autosome. The same 50% frequency would also hold true for the X-chromosome, except with the additional condition that a father would always pass the allele on to his daughters and never to his sons. Our observations of the sequenced progeny of heteroplasmic fathers in both the current and the previous manuscript [24] shows that 5 out of the 8 children that were sequenced received mtDNA from their father, which is consistent with the 50:50 transmission pattern expected for a Mega-NUMT. However, in every case where we have sequenced whole mtDNA from an individual whose mother carried two populations of mtDNA (17 individuals in total), that individual also inherited both of their mother’s mtDNA haplotypes. This includes 4 individuals from our previous publication [24], as well as 8 individuals from Family D (**Fig. 2**), 2 individuals from Family E (**Fig. 3**), and 3 individuals from Family F (**Fig. 4**). This is, of course, a classic pattern of strictly maternal transmission, which is exactly what one would expect if both mtDNA populations are genuine mtDNA. However, the chance of this occurring if these transmission events are due to Mega-NUMTs is 0.5^17, or about 0.00000763 (in other words, about 0.000763%). Among the eight maternally related, mixed haplotype individuals described recently by Parson et al. [48], only a single individual failed to inherit both haplotypes from their heteroplasmic mother (personal communication). This could be explained by one of the two haplotypes being lost due to the maternal mtDNA bottleneck. However, mitochondrial DNA FISH showed the mega-copy mtDNA to be present on chromosome 14. If this is the case, it does add a great deal of mystery as to the exact nature of these Mega-NUMTs, their underlying genetics and their biological consequences.

In summary, biparental mtDNA inheritance remains a truly unique and remarkable genetic phenomenon, and it raises many questions about the exact evolutionary and biological purpose of a strictly maternal mtDNA inheritance process. We may, however, be able to make some informed speculations as to the mechanism by which paternal transmission occurs. For instance, in species where paternal mitochondria are introduced into the egg they are rapidly destroyed [51]. One model proposes that the autosomal locus responsible for biparental inheritance is somehow disrupting this process, and that it is operating on the paternal side rather than the maternal side. Answering these questions may be of particular importance in the field of reproductive medicine, since the exclusive maternal inheritance of mtDNA creates a significant reproductive burden for women who are afflicted with deleterious mtDNA variants. While interventions such as oocyte spindle transfer have recently shown promising results in preventing the transmission of disease-causing mtDNA variants from mother to child, there are [52] numerous ethical and financial hurdles to be overcome with these techniques. The phenomenon of paternal mtDNA transmission, if properly explained and understood, may present an opportunity to develop alternate methodologies in this area. The ability to transmit healthy mtDNA from the father would be an excellent means of rescuing deleterious mtDNA variants transmitted from the mother, but only if the full genetic consequences of paternal mtDNA transmission are properly understood and sufficiently benign. To that end, the present results represent a significant step toward achieving a better understanding and development of paternal transmission of mtDNA as a practical tool.

## Acknowledgements

We thank the patients and their families who participated in this study, as well as Dr. Zitao Liu at New Hope, New York for helping with sperm collection. We would also like to thank the DNA Core facilities where next-generation sequencing was performed. This work was supported in part by the Cincinnati Children’s Hospital Research Foundation and NICHD grant, 1R01HD092989-01A1 to TH.

## Author contributions

JZ and TH conceived the idea for this study; JS, WZ, SL, SMC, JB, and MB performed experiments; JS, SL, ESS, XW, L-J W, JZ and TH analyzed data; Y-H C, W-L H, P-C F, N-C L, and TH evaluated patients and their families, and JS, ESS and TH wrote the manuscript. The project was designed and supervised by TH. This project was supported by the Cincinnati Children’s Hospital Research Foundation and NICHD grant, 1R01HD092989-01A1, to TH.

## EXPERIMENTAL PROCEDURES

### Patients

Four unrelated, multi-generation families with a high number and level of mtDNA heteroplasmy events were examined. The probands of Family A, Family D and Family F were evaluated at the MitoClinic at the Human Genetics Division of Cincinnati Children’s Hospital Medical Center (CCHMC); for Family A, some of these results were reported previously [24]. These families were subsequently referred to our Mitochondrial Diagnostic Laboratory. The proband of Family E was evaluated at National Taiwan University Hospital (NTUH) and Genetic testing for Family E was performed at the Baylor Genetics Laboratory before being referred to CCHMC for further evaluation. Details regarding the medical history of Family A have been described previously [24]. Additional details regarding Family D, Family E, and Family F are provided in the “Results” section. Written informed consent was obtained from all participating family members. This study was approved by the Institutional Review Board of CCHMC (Approval Study ID: 2013-7868), and all methods were performed in accordance with the relevant guidelines and regulations.

### Whole mitochondrial genome sequencing analysis

Whole mitochondrial genome sequencing was performed independently for each family at two different CLIA-accredited laboratories, CCHMC (**Method 1**) and Baylor Genetics (**Method 2**). Briefly, genomic DNA was isolated from various tissue samples collected from patients and their family members with a Gentra DNA extraction kit (Qiagen, Hilden, Germany), according to the manufacturer’s instructions. Entire mtDNA molecules were subsequently amplified in long-range PCR reactions as previously described [25–28]. For these reactions, 100 ng total genomic DNA were used as the templates in individual 50-μL reactions.

For **Method 1** (used for Family D), two sets of primers were used to specifically recognize genuine mtDNA: F-2120 (GGACACTAGGAAAAAACCTTGTAGAGAGAG) and R-2119 (AAAGAGCTGTTCCTCTTTGGACTAACA), and F-2662 (ACTTTTAACCAGTGAAATTGACCTGCC) and R-2661 (AAGAGACAGCTGAACCCTCGTGG). PCR amplification was performed with TaKaRa LA Taq Hot Start polymerase (TaKaRa Biotechnology, Kyoto, Japan) under conditions of: 94 °C for 1 min; 30 cycles of 98 °C for 10 s and 68 °C for 16 min; 72 °C for 10 min; and a final hold at 4 °C. The two resulting PCR products were mixed in equimolar amounts and each sample was barcoded according to the Nextera XT library preparation protocol (Illumina, San Diego, CA, USA). Sequencing was performed on the Illumina MiSeq platform (DNA Core Facility, CCHMC) and data were analyzed with NextGENe software (SoftGenetics, State College, PA, USA). Briefly, sequence reads ranging from 100–200 bps in length were quality filtered and processed with NextGENe software and an algorithm similar to BLAT. The sequence error correction feature (condensation) was performed to reduce false positive variants and to produce a sample consensus sequence and variant calls. Alignment without sequence condensation was performed to calculate the percentage of the mitochondrial genome with depth of coverage of 1000×. Quality FASTQ reads were quality filtered and converted to a FASTA format. Filtered reads were then aligned to the human mitochondrial sequence reference, NC_012920.1, followed by variant calling. Variant heteroplasmy was calculated with NextGENe software as follows: base heteroplasmy (mutant allele frequency %) = mutant allele (forward + reverse) / total coverage of all alleles C, G, T, A (forward + reverse) * 100. Clinical significance of the variants identified and the predicted haplogroups were analyzed with MitoMaster software (http://www.mitomap.org/MITOMASTER/WebHome) [25–27].

For **Method 2** (used for Family E), the forward and reverse primers used for long-range PCR included: mt16426F (CCGCACAAGAGTGCTACTCTCCTC) and mt16425R (GATATTGATTTCACGGAGGATGGTG). PCR amplification was performed with TaKaRa LA Taq Hot Start polymerase (TaKaRa Biotechnology) under conditions of: 95 °C for 2 min; 30 cycles of 95 °C for 20 s and 68 °C for 18 min; 68 °C for 20 min; and a final hold at 4 °C [28]. The amplified PCR products were fragmented to 200 bp, purified with AMPure XP beads (-), and subjected to end repair, 3’-adenylation, and Illumina InPE adapter ligation. The DNA samples were then enriched by PCR with Herculase II polymerase (Agilent Technologies, Santa Clara, CA, USA). Briefly, twelve indexed DNA libraries were pooled together in equimolar ratios. Each pooled library was sequenced in a single lane of one flow cell on an Illumina HiSeq2000 instrument with 76- or 100-bp paired-end or single-end read chemistry. After demultiplexing the reads with CASAVA software (Illumina), the reads belonging to the same index were filtered to remove any reads with a median quality score < 25 by using NextGENe software (SoftGenetics). Reads with one unknown assigned base call were also removed. The data were then processed and variant calls were made with two different heteroplasmy cutoff values, 20% and 1%. The aligned reads were subsequently examined with NextGENe Viewer [28]. To ensure that every indexed sample was scored correctly during pool sequencing, an internal identity control system (InQC) was designed and incorporated into the analysis of each sample. Briefly, fourteen nuclear DNA regions containing polymorphic markers were amplified in a multiplex PCR reaction. The resulting DNA fragments were mixed with the long-range PCR-enriched mtDNA fragments from the same individual prior to indexing and massive parallel sequencing. In parallel, genotyping was performed for the 14 nuclear markers in the same sample with TaqMan assays, or Sanger sequencing was performed independently. The genotyping results obtained from this parallel testing were compared to those generated with parallel sequencing to verify sample identity. To ensure that the quantification of heteroplasmy by massively parallel sequencing was reliable, a set of cloned synthetic 150-bp control DNAs was spiked into each indexed sample as an external identity control system (ExQC).

### Single-cell sequencing analysis

For the sequencing analysis of single-cell fibroblast samples, primary dermal fibroblasts obtained by needle biopsy were plated at a low density to allow distinct, clonal colonies to form. Once these colonies were established, the individual colonies were briefly trypsinized and isolated using a micropipette. DNA was then extracted from each sample, and next-generation sequencing (NGS)-based whole mitochondrial genome sequencing analysis was performed for each of the DNA samples according to “**Method 1**” (described above).

To analyze single-sperm mtDNA sequences, an aliquot of frozen sperm from Patient II-4 (Family A) was thawed and washed once in high calcium HTF media (CytoSpring LLC, Mountain View, CA, USA). The sperm sample was then reconstituted in high calcium HTF plus 10% PVP media (Sigma-Aldrich, St. Louis, MO, USA). Individual sperm were sorted into separate droplets of HTF with a piezo micromanipulator (Eppendorf, Hamburg, Germany) and placed into separate PCR tubes for long-term storage at −80 °C. A total of 44 single-sperm samples were selected to undergo whole genome amplification (WGA) with the REPLI-g Single Cell Kit (Qiagen, Hilden, Germany), according to the manufacturer’s protocol. The products obtained were subsequently subjected to semi-nested PCR with a LA Taq Hot Start polymerase kit (TaKaRa Biotechnology). The first round of amplification was performed with primers, hmtF2-9611 (TCCCACTCCTAAACACATCC) and hmtR2-626 (TTTATGGGGTGATGTGAGCC). The second round of amplification was conducted with primers, mt16426F (CCGCACAAGAGTGCTACTCTCCTC) and hmtR2-626 (TTTATGGGGTGATGTGAGCC). PCR conditions for the first round of amplification included: 95 °C for 2 min; 35 cycles of 95 °C for 20 s and 68 °C for 16 min; 68 °C for 20 min; and a final hold at 4 °C. For the second round of amplification, the PCR conditions were: 95 °C for 2 min; 35 cycles of 95 °C for 20 s and 68 °C for 1 min; 68 °C for 10 min; and a final hold at 4 °C. Eleven reactions produced bands visible in gel electrophoresis. Ten of these PCR products were purified with ExoSapIT (Thermo Fisher Scientific, Waltham, MA, USA) and were then submitted for Sanger sequencing with primer, mtDNA-207R (CACACTTTAGTAAGTATGTTCGCC). The sequencing results were compared against expected sequences for haplogroups R0a1 and H1a1 to estimate the level and type of heteroplasmy in each sample.

