## Supplemental Material for "Heteroplasmy variability in individuals with biparentally inherited mitochondrial DNA"

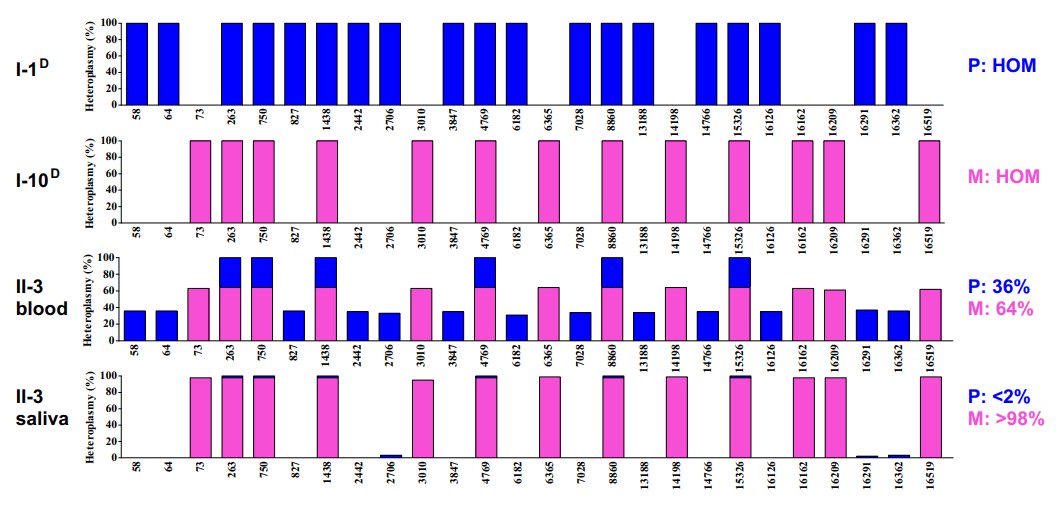

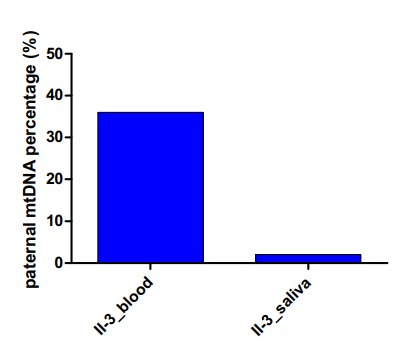

**Figure S1. Tissue-related variations in heteroplasmy levels for maternal (H1a1) and paternal (R0a1) haplogroups in Patient II-3 of Family A.** PCR-NGS detected variable heteroplasmy levels in blood and saliva samples obtained from Patient II-3.

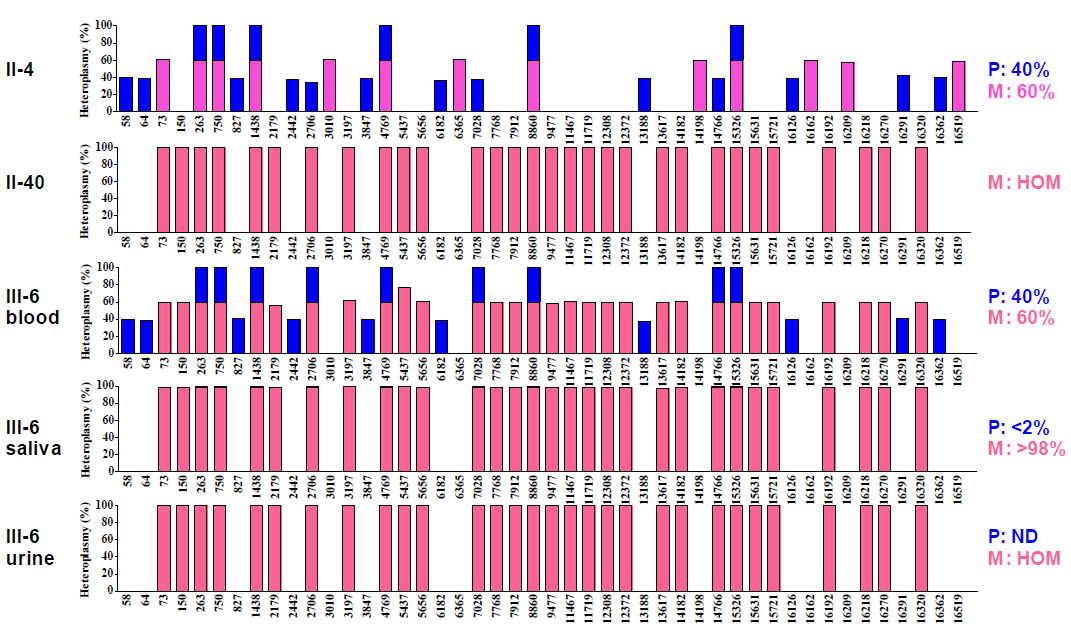

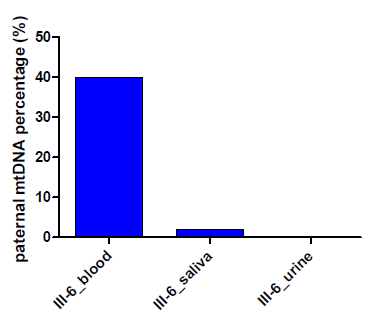

**Figure S2. Tissue-related variations in heteroplasmy levels for maternal (U5b1d1c) and paternal (R0a1) haplogroups in Patient III-6 of Family A.** PCR-NGS detected variable heteroplasmy levels in blood, saliva, and urine samples obtained from Patient III-6.
